# Identification of S1PR4 as an immune modulator for favourable prognosis in HNSCC through unbiased machine learning

**DOI:** 10.1101/2022.12.20.521217

**Authors:** Chenshen Huang, Fengshuo Zhu, Ning Wang, Qi Huang

**Affiliations:** Department of Gastrointestinal Surgery, Fujian Provincial Hospital, Shengli Clinical Medical College of Fujian Medical University, Fuzhou, China; Department of General Surgery, Tongji Hospital, Tongji University School of Medicine, Shanghai, China; Department of Oral Maxillofacial-Head and Neck Oncology, Shanghai Ninth People’s Hospital, College of Stomatology, Shanghai, China. Jiao Tong University School of Medicine, National Clinical Research Center for Oral Disease, Shanghai Key Laboratory of Stomatology and Shanghai Research Institute of Stomatology, Shanghai, China; Department of Hepatobiliary and Pancreatic Surgery, Huzhou Central Hospital, Huzhou, China

**Keywords:** S1PR4, CD8 T cell, Immune modulator, HNSCC, Unbiased machine learning

## Abstract

As the largest family of membrane proteins, G protein-coupled receptors (GPCRs) are the most prominent family of pharmacological targets. However, only a few GPCRs have been well-defined in terms of their physiological and pathological functions. Thus, an efficient way to identify key GPCRs involved in tumour formation is urgently needed. In this study, patients with head and neck squamous cell carcinoma (HNSCC) were classified into two different subtypes based on the characteristics of GPCRs using an unbiased machine learning method. Notably, these two subtypes showed significant differences in prognosis, gene expression, and immune microenvironment, especially in the infiltration of CD8^+^ T cells. Based on these differences, we screened three potential key regulators (S1PR4, S1PR5, and GPR87) from all GPCRs and constructed a prognostic nomogram for patients with HNSCC. We identified S1PR4 as the key GPCR in determining the two subtypes for positive correlation with the proportion and cytotoxicity of CD8^+^ T cells in HNSCC and was mainly expressed in a subset of CX3CR1^+^CD8^+^ T cells. We also demonstrated that S1PR4 is an immune modulator for the favourable prognosis of HNSCC patients. We found that S1PR4 was highly expressed in CD8^+^ T cells from the tumours of HNSCC patients, which was significantly associated with better prognosis, and S1PR4 expression was accompanied by higher T cell cytotoxic marker expression (IFNG and GZMB). Notably, S1PR4 co-localised with CX3CR1, which has been identified as the most cytotoxic marker of CD8^+^ T cells. Furthermore, S1PR4 upregulation could significantly increase T cell function in CAR-T cell therapy, indicating its great potential in cancer immunotherapy. Therefore, these results identified S1PR4 as a key indicator of cytotoxicity and the proportion of tumour-infiltrating CD8^+^ T cells and confirmed the prognostic value of S1PR4 in HNSCC. Our findings contribute to the knowledge of S1PR4 in anti-tumour immunity, providing a potential GPCR-targeted therapeutic option for future HNSCC treatment.

## Introduction

G protein-coupled receptors (GPCRs), which have a seven-transmembrane structure, comprise the largest family of cell-surface signalling proteins. Thus far, the GPCR family has been regarded as the most prominent membrane protein family of pharmacological targets for the treatment of human diseases. For instance, many currently marketed anti-tumour drugs are GPCRs[1, 2]. Meanwhile, GPCRs can be activated by various important active mediators in the immune or inflammatory response, such as chemokines, S1P, and lysophosphatidic acid (LPA), which play key roles in cell proliferation, transformation, angiogenesis, and metastasis, as well as in inflammation-associated cancer[1]. However, some GPCRs, such as OX1R, have been shown to promote robust apoptosis in various cancer cells[3]. To date, more than 800 membrane proteins have been identified in the human genome as GPCRs, and the role of numerous GPCRs in tumour formation and progression remains unclear[4]. Accordingly, an improved understanding of GPCRs’ involvement in tumour formation and progression may contribute to the development of a new generation of anti-tumour drugs.

Owing to the heterogeneity of tumours, different types of tumours often have their characteristics. Therefore, therapeutic targets developed for specific tumour types often have better clinical effects. In our study, we focused on head and neck squamous cell carcinoma (HNSCC), which is the sixth most common malignancy worldwide, accounting for approximately 380,000 deaths and 600,000 new cases annually[5, 6]. Currently, with limited treatment options, the prognosis of HNSCC patients remains poor. Therefore, a more effective treatment strategy for HNSCC is still warranted[7-9]. The tumour microenvironment (TME) plays a key role in HNSCC progression. The TME of HNSCC has typical features, such as amplification of circulating Tregs, high levels of TAMs, and expression of high levels of TGFβ and VEGF, which result in Treg cell activation, NK cell suppression, suppression of DC maturation, and T-lymphocyte inactivation and dysregulation[10, 11]. High levels of cytotoxic CD8^+^ T lymphocytes are a favourable prognostic factor for oropharyngeal cancer[12]. Most importantly, some GPCRs, such as CX3CR1, can improve the cytotoxicity of CD8^+^ T cells[13, 14]. Therefore, this study aimed to explore the key GPCRs involved in HNSCC development, providing a potential GPCR-targeted therapeutic option for HNSCC treatment.

Recently, multiple machine learning methods, including non-negative matrix factorisation (NMF), least absolute shrinkage and selection operator (LASSO), and Random Forest (RF), have been widely used to explore key biomarkers in patients with tumours[15]. With the methods mentioned above, previous studies could classify tumour patients into different molecular subtypes and could further identify the potential key biomarkers by comparing the classified subtypes. However, in previous studies, the gene list of specific biological functions or pathways (for example ferroptosis, N6-methyladenosine, or hypoxia) has often been determined in advance. Subsequently, the molecular subtypes of the patients were further classified based on these preselected genes. Therefore, at the beginning of these studies, the biological differences were limited to specific biological functions or pathways to which the potential key biomarkers were anticipated to be related. In our study, we decided not to limit the biological function or gene pathways, and not to anticipate biological differences between molecular subtypes. Thus, we classified the patients into different molecular subtypes based on GPCRs, which were determined by their structures rather than specific biological functions, and were regarded as the most prominent family of pharmacological targets.

Here, we first classified HNSCC patients into two different GPCR-based molecular subtypes, only from a statistical perspective. We then compared the differences between the two subtypes from clinical and biological perspectives and identified potential key GPCRs in determining these two subtypes. We also explored the prognostic value of key GPCRs and their associations with the immune microenvironment. We validated our findings using an independent cohort of 59 HNSCC patients from our institution using immunohistochemistry (IHC), immunofluorescence (IF), and Kaplan-Meier survival analysis (K-M analysis). We evaluated the immunotherapeutic potential of S1PR4 using CAR-T cells and the S1PR4 inhibitor/agonist. We found that S1PR4 could be a key regulator of the anti-tumour response of CAR-T cells.

## Materials and Methods

### Ethics

In this study, an independent cohort was used for validation, which included 59 HNSCC patients with a median follow-up time of 4.17 years. All patients provided written informed consent before surgery. This study was reviewed and approved by Shanghai Ninth People’s Hospital, Shanghai, China. The reference number for ethical approval was SH9H-2022-T189-1.

### Data Collection

We retrieved the gene expression profiles of the TCGA database using UCSC Xena (http://xena.ucsc.edu/). Htseq-count profiles and Fragments per Kilobase per Million mapped reads (FPKM) profiles of the TCGA-HNSCC cohort were downloaded, including 500 primary HNSCC, two metastatic HNSCC sample, and 44 solid normal tissue samples. Subsequently, the Htseq-count and FPKM profiles of the primary HNSCC samples were retrieved. The survival data and clinical demographic information of the primary HNSCC patients was also acquired using UCSC Xena.

### NMF Clustering

A total of 500 patients with HNSCC were clustered into different subtypes using the NMF method. The R package “NMF” was used to perform NMF clustering with the “brunet” criterion and 100 interactions[16, 17]. The number of clusters, *k*, was set from 2 to 10. The minimum number of patients in each subtype was set to 10. The NMF rank survey was performed to determine the optimal cluster numbers. The stability of the clusters was reflected by the cophenetic correlation coefficient. According to literature investigation, the cophenetic correlation coefficient is currently the most common approach for determining the optimal values of rank *k*[16]. The values of rank *k* were selected where the cophenetic correlation coefficient started decreasing or was found to decrease most sharply. In this study, both conditions occurred at rank *k* = 2. Therefore, we selected rank *k* = 2 as the optimal value, clustering the samples into two subtypes.

### Identification of the Differentially Expressed Genes

We used the R package “edgeR”[18] to identify the differentially expressed genes (DEGs) between molecular subtypes. For the screening of DEGs, statistical significance was set at FDR p < 0.05. And the log2 (Fold Change) > 1 or < -1 was defined as either upregulated or downregulated, respectively. Heatmap and volcano plot were used to visualise DEGs based on the R package “pheatmap” and the R package “EnhanceVolcano”.

### Gene Set Enrichment Analysis

The algorithm of Gene Set Enrichment Analysis (GSEA) was used to explore the differences in pathway expression between molecular subtypes[19]. Gene sets of the HALLMARK, KEGG, and GO pathways were identified using the Molecular Signatures Database (http://www.gsea-msigdb.org/). The above gene sets were obtained using the R package “msigdbr”. The R packages “clusterProfiler”[20, 21] and “enrichplot” were used to perform the GSEA algorithm between the molecular subtypes. Statistical significance was set at p.adjust < 0.05.

### Estimation of Immune Infiltrating Microenvironment

ESTIMATE scores were employed for evaluating the immune-infiltrating microenvironment for each sample. Subsequently, the MCPcounter algorithm[22] was used to estimate the proportion of infiltrating immune cells in HNSCC samples. To minimise the bias of using a single algorithm, four other algorithms (CIBERSORT[23], CIBERSORT-ABS[24], QUANTISEQ[25], and TIMER[26]) were further performed for validation. The R package “limma”[27] was used to identify immune cells with significant changes in proportion between molecular subtypes. The R package “factoextra” was used for Principal Component Analysis (PCA).

### Construction of Prediction Nomogram

The R package “glmnet”[28] was used to perform LASSO regression, ensuring that there was no overfitting in the model. Subsequently, the reduced multi-cox model was constructed using the R package “Survival”. Based on the model, the prediction nomogram was visualised using the R package “rms”. Using the R package “nomogramEx”, the acceptable accuracy of the prediction nomogram was validated using calibration curves of 3-year-survival and 5-year-survival.

### Integrative Analysis for Key GPCR Exploration

To explore the key GPCRs, we proposed a strategy of “classification before analysis”. We first clustered HNSCC patients into different subtypes based on genes with a similar structure, instead of genes related to specific biological functions/pathways. Then, from multiple perspectives, we confirmed the differences in the characteristics between the subtypes. Next, we analysed the clustering principle and identified GPCRs that might be essential in the clustering procedure. Eventually, among these GPCRs, we searched for the key GPCR that was closely correlated with the featured characteristics.

The detailed steps were as follows: **1)** The list of currently known GPCRs was downloaded from the GPCR NaVa database (http://nava.liacs.nl)[29], and gene expression of the recorded GPCRs was retrieved from the Htseq-count profiles of primary HNSCC patients. **2)** These GPCRs were filtered by their average expression. The GPCRs with an average expression > 1 were further filtered using univariate Cox analysis. **3)** Based on the retained GPCRs, we performed NMF to cluster HNSCC patients into different subtypes. The K-M analysis was used to confirm the difference in prognosis between subtypes. The differences in characteristics (gene expression, pathway expression, tumour microenvironment, and immune cells) between the subtypes were also investigated using the methods, which had been described in the previous sections. **4)** After confirming significant differences between subtypes, we analysed the principle of clustering. As the subtypes were clustered based on the featured GPCRs in step 3, these GPCRs were further investigated. We used LASSO regression and reduced Multi-Cox regression to screen the potential key GPCRs. **5)** With the R package “randomForest”, we employed the RF method to compare the importance of the potential key GPCRs. The prognostic value of these GPCRs was also compared using K-M analysis. The cutoffs were determined using either maximally selected rank statistics or median gene expression. The Pearson correlation analysis of these GPCRs and CD8^+^ T-cell-related markers (CD8A, CD8B, IFNG, and GZMB) was also performed.

### Multi-database Analysis

We acquired and analysed the single-cell RNA-seq profiles of T cells in pan-cancer using the scDVA data portal (http://cancer-pku.cn:3838/PanC_T/) to identify the cell subset in which the key GPCR was expressed. The meta-cluster of CD8^+^ T cells, proposed by Zheng et al.[13], was visualised using the default settings of the scDVA data portal. The distribution of the key GPCR was analysed and visualised by the “Embedding” function of the scDVA data portal.

In this study, we used multiple databases to validate our results, preventing the bias of using only a single database. The prognostic value of the key GPCR was validated by several databases, including the Timer database[26, 30, 31], LinkedOmics database[32], and HumanProteinAtlas database (https://www.proteinatlas.org/). Correlations between the key GPCR and several marker genes were validated using the Gepia database[33].

### IHC Staining

IHC staining was performed to detect IFNG and GZMB protein expression in our independent HNSCC cohort. The 59 paraffin-embedded HNSCC samples were incubated overnight at 4 °C with primary antibodies against IFNG (Abcam, ab231036) and GZMB (Abcam, ab134933). The slides were then washed thrice and incubated with the secondary antibody for 1 h at room temperature. DAB (Service Bio, G1211) was used to stain the slides, and haematoxylin was used for nuclear staining. IHC staining results were detected using a microscope from the ServiceBio platform (ServiceBio, Wuhan, China).

### IF Staining

Paraffin-embedded tissues from 59 HNSCC samples from our independent cohort were used for IF staining. Primary antibodies, including S1PR4 (Affinity, DF4872), CD8 (ServiceBio, GB13068), CX3CR1 (ServiceBio, GB11861), IFNG (Abcam, ab231036), and GZMB (Abcam, ab134933), were used to detect the target proteins. DAPI (ServiceBio, G1012) was used for counterstaining of the nuclei. IF staining results were detected using a microscope from the ServieBio platform (ServiceBio, Wuhan, China).

### Preparation of CAR-T Cells

Peripheral blood for CAR-T cells was obtained from healthy volunteers (n = 3 or more), and written informed consent was obtained from all participants. All blood samples were collected and handled according to ethical and safety procedures approved by the Clinical Ethics Committee of the First Affiliated Hospital, College of Medicine, Zhejiang University (IIT20210001C-R1 for human subjects). Human CD4^+^ and CD8^+^ T cells were purified from PBMCs using human CD4 (Miltenyi, 130-045-101) and CD8A magnetic microbeads (Miltenyi, 130-045-201). After two days of activation with CD3 and CD28, T cells were used for the experiment. We constructed a second-generation CAR model. The first nucleic acid sequence of the CAR comprised the anti-human PSMA-scfv (J591) linked in-frame to the hinge and transmembrane regions of the human CD8α chain and intracellular human 4-1BB (CD137) and CD3ζ signalling domains, which were transferred to the plasmid-empty vector (lentiviral transfer vector pELPS) using XbaI and Sal? double enzymatic digestion. For lentivirus production, 293 T cells were used to produce lentiviruses carrying CAR. Subsequently, 293 T cells were cultured to a degree of 80%-90% and co-transfected with the three-plasmid system (packaging vectors psPAX2, envelope plasmid pMD2.G, and CAR plasmid) using polyethyleneimine (Polysciences, PEI MAX 40000). At 6-8 h of post-transfection, the virus-containing supernatant was harvested and concentrated. Viral supernatants generated were serially diluted, and then T cells were infected with the lentivirus, which was mixed evenly with Polybrene at MOI = 10. CAR expression was analysed after two or three days, and CAR-T cells were used for further experiments.

### Cell preparation and culture

The prostate cancer cell line (PC3) was obtained from the American Type Culture Collection (ATCC). PC3 cells were cultured in DMEM (Gibco) supplemented with 10% FBS and 1% P/S (100 U/mL penicillin, 100 mg/mL streptomycin). Prostate-specific membrane antigen (PSMA) was overexpressed in PC3 cells using lentivirus. The cells were then screened with puromycin dihydrochloride (2-4 μg/ml; BBI Life Science, F118BA0026) to stablely overexpress PSMA and luciferase. PC3 cells were cultured at 37 °C and 5% CO_2_ and were regularly tested for mycoplasma-free status.

### Cytotoxicity assay

The cytotoxicity of CAR-T cells was determined using a luciferase-based killing assay. PSMA-CAR-T cells were pretreated with DMSO, S1PR4-inhibitor (CYM50358, MedChemExpress, 100 nM) and S1PR4-agonist (CYM50308, MedChemExpress, 100 nM) for 2 h. 2×10^4^ PC3-PSMA^+^ target tumour cells expressing firefly luciferase were co-cultured with PSMA-CAR-T cells at an effector-to-target ratio of 4:1 in triplicates in 96-well plates (Corning) in a total volume of 100 μl of cell media. After 12 h, 10 μL of fluorescein substrate (Invitrogen) was added to each well and luminescence was measured using a microplate reader (BMG LABTECH SPECTROstarNano, 562 nm) and IVIS instrument (IVIS Lumina III). Percent lysis was determined using a standard curve by linear regression of luminescence against the viable number of PC3-PSMA^+^ cells.

### Cytokine detection

PSMA-CAR-T cells were pretreated with DMSO, S1PR4-inhibitor (CYM50358, MedChemExpress, 100 nM) and S1PR4-agonist (CYM50308, MedChemExpress, 100 nM) for 2 h. PSMA-CAR-T cells were co-cultured without PC3-PSMA^+^ target tumour cells (negative control) or with 1 × 10^6^ PC3-PSMA^+^ target tumour cells at an effector-to-target ratio (2:1) in triplicates in 6-well plates (Corning) in a total volume of 2 ml of cell media. After 16 h, PSMA-CAR-T cells were collected and GZMB was measured using flow cytometry. All samples were run on an LSRFortessa (BD Pharmingen) and analysed using FlowJo software (Tree Star). The concentration of IFNG in the culture medium was measured by ELISA (Invitrogen).

### Statistical Analysis

Statistical significance was set at adjusted p < 0.05 or p < 0.05. The correlation threshold was set at r < -0.2 or r > 0.2 in Pearson or Spearman correlation analysis. The Shapiro-Wilk normality test was applied for checking the variable normality. For the variables which distributed non-normally, we applied Wilcoxon rank test for comparisons between two independent groups. For normally distributed variables, we applied Student’s *t*-test for comparisons of two independent groups. In the K-M analysis, the optimal cutoffs were determined by the median gene expression, or by the maximally selected rank statistics. When the K-M analysis was performed in TCGA-HNSCC cohort, which had a relatively larger number of patients, both methods of cutoff determination were used. When the K-M analysis was performed in our independent cohort, which included only 59 patients, we chose to use the median gene expressions as cutoffs. The method of maximally selected rank statistics was performed using the R package “survminer”. All statistical analyses in this study were performed using R software (Version 3.5.1, http://www.r-project.org/).

## Results

### Identification of GPCR-Based Molecular Subtypes *via* NMF Method

NMF is an effective machine learning method, which has been widely used in computational biology for molecular pattern discovery, class comparison, and class prediction[34, 35]. Here, we applied the NMF method to explore the GPCR-based molecular subtypes in the TCGA-HNSCC cohort.

First, we obtained a list of currently known GPCRs from the GPCR NaVa database (http://nava.liacs.nl). Ninety GPCRs with an average expression > 1 in the TCGA-HNSCC cohort were filtered for further analysis. Further filtering procedures were performed using univariate Cox analysis; 17 GPCRs with significant prognostic value for 5-year survival were retained for NMF clustering (p < 0.05, **Figure 1A**). We performed the NMF rank survey (**Figure 1B** and **Figure S1**) for clustering prediction, and noticed that the magnitude of the cophenetic correlation coefficient began to fall sharply at rank *k* = 2 (**Figure 1B**). According to the sharp decrease, substantially less stability is achieved with more than two clusters. However, the consensus maps of NMF clustering (rank *k* was set from 2 to 10) also indicated that the clustering results were better when rank *k* = 2 (**Figure S1**). Therefore, we decided to select rank *k* = 2 as the optimal value and clustered HNSCC patients into two GPCR-based molecular subtypes (C1 and C2, **Figure 1C**). The gene expression of 17 featured GPCRs used in NMF clustering was visualised using a heatmap (**Figure 1D**) and a volcano plot (**Figure 1E**). After NMF clustering, there were 363 patients in subtype C1, and 137 patients in subtype C2.

**Figure 1.**
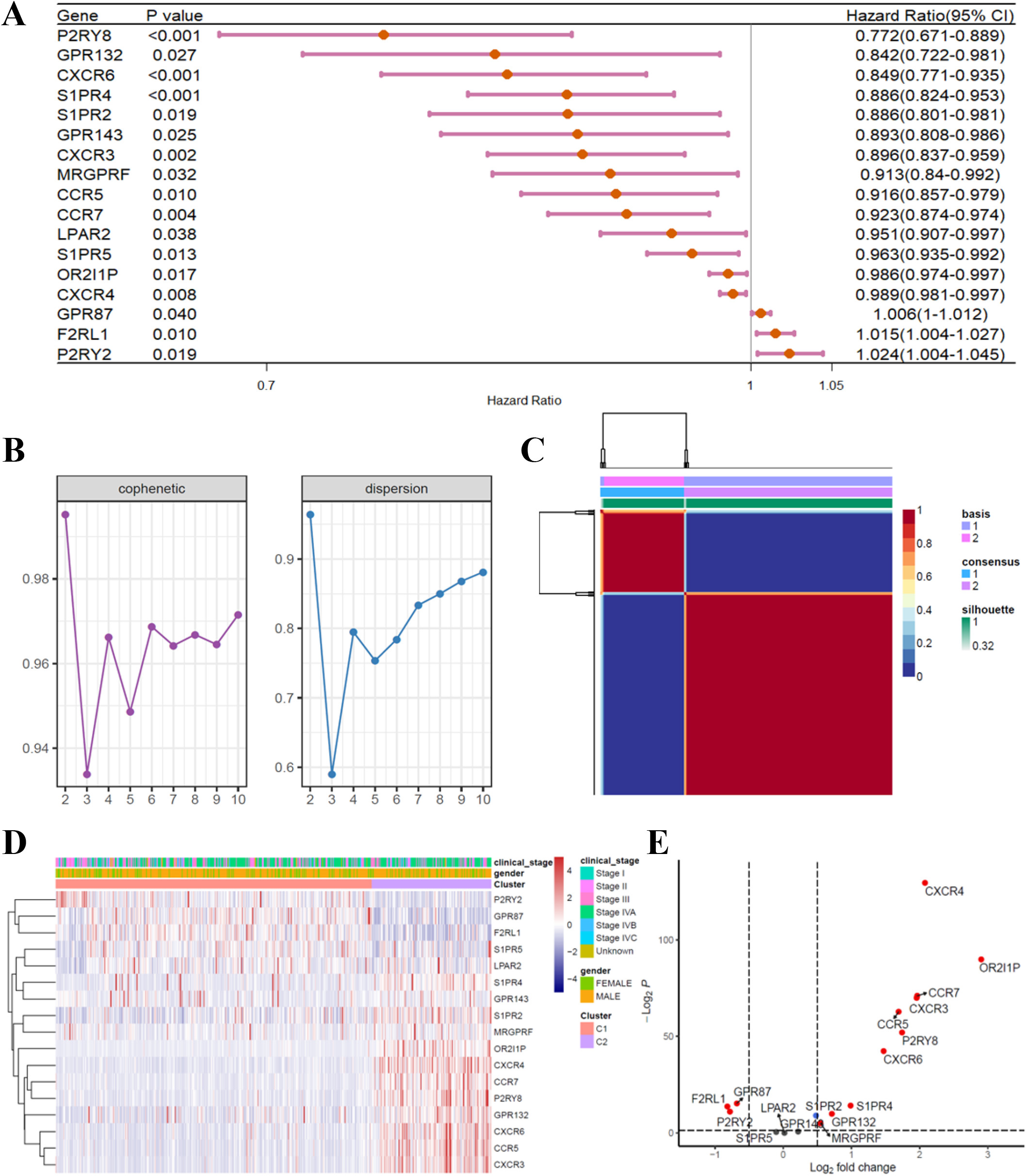
Identification of GPCR-Based Molecular Subtypes via NMF Method. **(A)**Filtered by univariate cox analysis, 17 GPCRs were found to have significant prognostic values of 5-year survival (p < 0.05), and were retained for further analysis. **(B)**The NMF rank survey was performed for clustering prediction (rank k was set as 2 to 10). The cophenetic correlation coefficient started decresing at rank k = 2, and was found to decreased most sharply at rank k = 2. Similarly, the sharp decrease of dispersion value was also found at rank k = 2. **(C)**Consensus map of NMF clustering (rank k = 2). The HNSCC patients were clustered into two GPCR-based molecular subtypes, including 363 patients in subtype C1, and 137 patients in subtype C2. **(D)**Gene expression of the 17 featured GPCRs was visualized through heatmap. **(E)**Volcano plot of the 17 featured GPCRs. The GPCRs which were differentially expressed between the subtypes were highlighted in red. *GPCR, G protein-coupled receptor; NMF, Non-negative Matrix Factorization; HNSCC, head and neck squamous cell carcinoma*.

### Differences in the Characteristics between GPCR-Based Molecular Subtypes

Because NMF is a machine learning method that does not consider biological significance, we further explored the differences between the identified subtypes (C1 and C2) from the perspective of biomedicine.

First, we compared the prognostic differences between the two subtypes. According to the K-M analysis, the overall survival (OS) time of HNSCC patients in subtype C1 was significantly shorter (p < 0.05, **Figure 2A**). Similarly, we also found that the progression free survival (PFS) time of subtype C1 tended to be shorter (p < 0.05, **Figure 2B**). Considering both the OS and PFS times, we found that the prognosis of patients in subtype C2 was significantly better.

**Figure 2.**
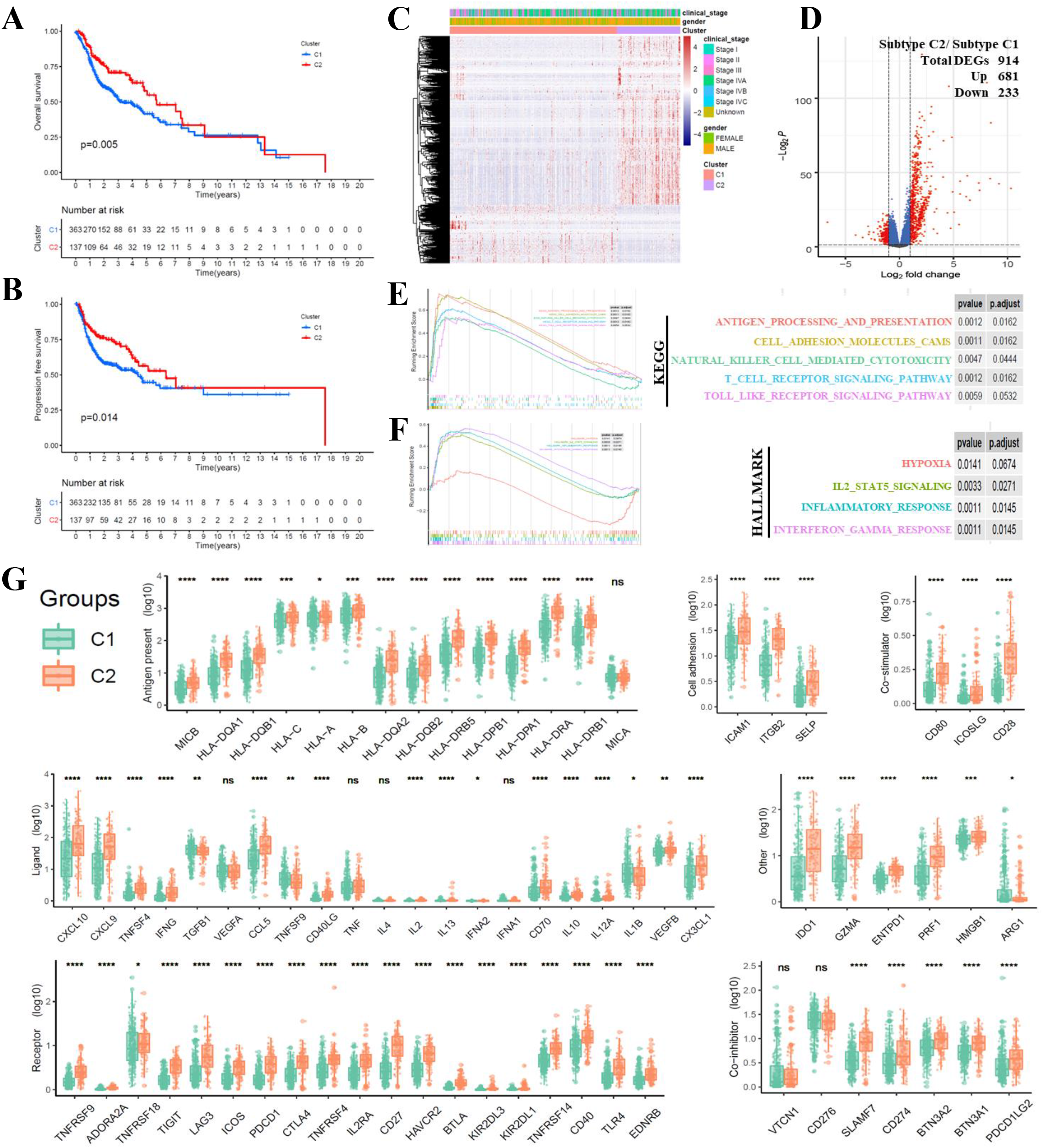
Differences of the Characteristics between GPCR-Based Molecular Subtypes. **(A)**Kaplan-Meier plot showed that the OS time of HNSCC patients in subtype C1 was significantly shorter (p < 0.05). Patients in Subtype C2 tended to have better prognosis of OS. **(B)**Kaplan-Meier plot showed that the PFS time of HNSCC patients in subtype C1 was significantly shorter (p < 0.05). Patients in Subtype C2 tended to have better prognosis of PFS. **(C, D)** A total of 914 genes were identified as DEGs between the subtypes. The gene expression of DEGs was visualized through heatmap **(C)**. The DEGs between subtype C1 and subtype C2 were highlighted in red **(D)**, including 681 and 233 upregulated and downregulated, respectively. **(E, F)** GSEA analysis was performed on KEGG pathways **(E)** and HALLMARK pathways **(F)**. The p < 0.05 was considered significant. **(G)** Seven classical series of genes were compared between the two subtypes. Scattered dots represented the relative gene expression values. *GPCR, G protein-coupled receptor; OS, overall survival; HNSCC, head and neck squamous cell carcinoma; PFS, progression free survival; DEG, differentially expressed gene; GSEA, Gene Set Enrichment Analysis. * p < 0*.*05; ** p < 0*.*01; *** p* *< 0*.*001; **** p < 0*.*0001; ns, not significant*.

We then explored the differences in gene and pathway expression between these two subtypes. Using the R package edgeR, 914 genes were identified as DEGs, including 681 and 233 upregulated and downregulated DEGs, respectively (**Figure 2C** and **Figure 2D**). To explore the differences in pathway expression, we performed the GSEA algorithm on KEGG and HALLMARK pathways. As shown in **Figure 2E** and **Figure 2F**, many immune-related pathways were found to be upregulated in the subtype C2 (p < 0.05). We also compared several classical series of genes related to antigen presentation, cell receptors, ligands, cell adhesion, co-inhibition, and co-stimulation between the two subtypes. Most of these genes were significantly upregulated in subtype C2 (**Figure 2G**). Therefore, we hypothesised that differences in the tumour immune microenvironment (TIME) might be one of the primary differences between the two subtypes.

### Cytotoxicity and Proportion of CD8^+^ T Cells were enhanced in Subtype C2

We then explored the differences in TIME between GPCR-based molecular subtypes. The ESTIMATE scores were important indicators of overall changes in TIME. We found that three immune-related ESTIMATE scores (ESTIMATEScores, ImmuneScores, and StromalScores) were significantly increased in subtype C2, whereas the tumour-related ESTIMATE score (TumorPurity) was relatively lower in subtype C2 (p < 0.001, **Figure 3A**). We also used the method proposed by Petitprez et al.[36] to estimate the gene signatures related to the functions of TIME and the expression of genes related to immune checkpoints. Genes associated with T cell activation and survival, Class I MHC, regulatory gene signatures, tertiary lymphoid structures (TLSs), and several immune checkpoint-related genes were all found to be higher in subtype C2 (**Figure 3B**). The above findings indicated that subtype C2 had an “immune and TLS high” characteristic, which was once described and shown to be correlated with better prognosis by Petitprez et al.[36] in sarcoma.

**Figure 3.**
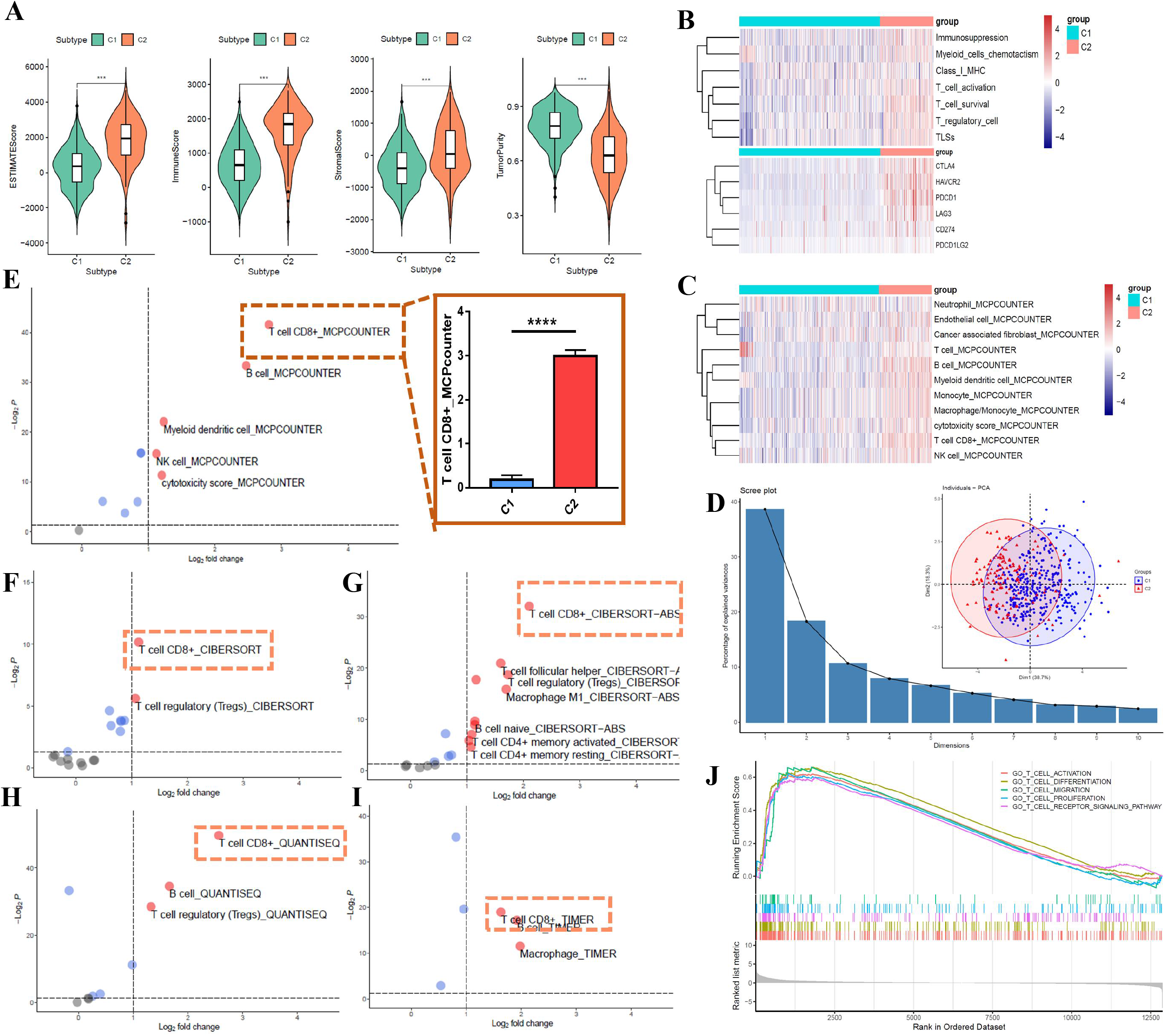
Cytotoxicity and Proportion of CD8^+^ T Cells were Upregulated in Subtype C2. **(A)**Violin plot showed the ESTIMATE scores of the two subtypes. Three immune-related ESTIMATE scores (ESTIMATEScores, ImmuneScores, and StromalScores) were significantly upregulated in subtype C2, while the tumour-related ESTIMATE score (TumorPurity) was relatively lower in subtype C2 (p < 0.05). **(B)**Heatmap for the expression of gene signatures, which were related to the functions of TIME or to the immune checkpoints. The gene signatures associated with T cell activation and survival, Class I MHC, regulatory gene signatures, TLSs, and several immune checkpoint-related genes were all found to be higher in subtype C2. **(C)**Heatmap for the proportion of TIICs, which were estimated by the MCPcounter algorithm. **(D)**PCA plot showed that the proportion of TIICs differed significantly between subtypes. **(E)**Volcano plot of the TIICs, which were estimated by the MCPcounter algorithm. The CD8^+^ T cells showed the largest value of fold-change, and the smallest value of p-value, indicating that among all the estimated TIICs, the largest difference in proportion was CD8^+^ T cells. The proportion of CD8^+^ T cells was significantly higher in subtype C2 (p < 0.0001). **(F-I)** Vocano plots of the TIICs, which were estimated by four algorithms, including CIBERSORT **(F)**, CIBERSORT-ABS **(G)**, QUANTISEQ **(H)**, and TIMER **(I)**. The CD8^+^ T cells were all found to have the largest value of fold-change, as well as the smallest value of p-value. **(J)** GSEA analysis of the GO pathways indicated that functional pathways, especially the cytotoxic function, of CD8^+^ T cells were significantly upregulated in subtype C2 (p < 0.05). *TIME, tumour immune microenvironment; TLS, tertiary lymphoid structure; TIIC, tumour-infiltrating immune cell. *** p < 0*.*001; **** p < 0*.*0001*.

After discovering the differences in the overall level of TIME, we further investigated the differences in tumour-infiltrating immune cell (TIIC) proportions between the two subtypes. To render the TIICs of each HNSCC patient comparable, we estimated the scaled proportion of TIICs using the MCPcounter algorithm (**Figure 3C**). The proportion of TIICs differed significantly between the subtypes (**Figures 3C** and **3D**). The volcano plot indicated that among all TIICs, the largest difference in proportion was CD8^+^ T cells (**Figure 3E**). Significant differences in CD8^+^ T cells were validated using four algorithms: CIBERSORT (**Figure 3F**), CIBERSORT-ABS (**Figure 3G**), QUANTISEQ (**Figure 3H**), and TIMER (**Figure 3I**). All the algorithms above confirmed that CD8^+^ T cells had the highest range of changes in proportion between the two subtypes. Therefore, the subtype C2 had a significantly higher proportion of CD8^+^ T cells. Moreover, GSEA analysis was performed to demonstrate differences in T cell functions. The functional pathways of CD8^+^ T cells were significantly upregulated in subtype C2 (p < 0.05, **Figure 3J**). In summary, both the proportion and cytotoxicity of CD8^+^ T cells were significantly elevated in the subtype C2, which was consistent with our previous finding that subtype C2 had a better prognosis.

### Identification of the Key GPCR S1PR4 in HNSCC

Next, we aimed to identify potential key GPCRs by analysing the principle of grouping two GPCR-based molecular subtypes. Considering that the subtypes were clustered based on GPCRs, we further investigated the 17 featured GPCRs used in the initial NMF clustering. First, the survival analysis indicated that all 17 GPCRs showed prognostic value for the OS of HNSCC patients (**Figure S2**). These GPCRs were further processed using the LASSO method (**Figures 4A** and **4B**). Based on the results of LASSO regression, five GPCRs (CXCR6, GPR87, MRGPRF, S1PR4, and S1PR5) were considered eligible and were further included in the reduced-Cox regression model (C-index = 0.61, p < 0.05, **Figure 4C**). The reduced-Cox model indicated that GPR87 (p = 0.048), S1PR4 (p = 0.030), and S1PR5 (p = 0.029) could effectively evaluate the prognosis of OS. More specifically, HNSCC patients with higher S1PR4 or S1PR5 expression tended to have a better prognosis for OS (hazard ratio < 1), whereas patients with higher GPR87 expression tended to have a poorer prognosis for OS (hazard ratio > 1). Moreover, we constructed a nomogram based on GPR87, S1PR4, and S1PR5 expressions (**Figure 4D**). The acceptable accuracy of this nomogram was validated by the calibration curves of the 3-year survival (**Figure 4E**) and 5-year survival (**Figure 4F**) of HNSCC patients.

**Figure 4.**
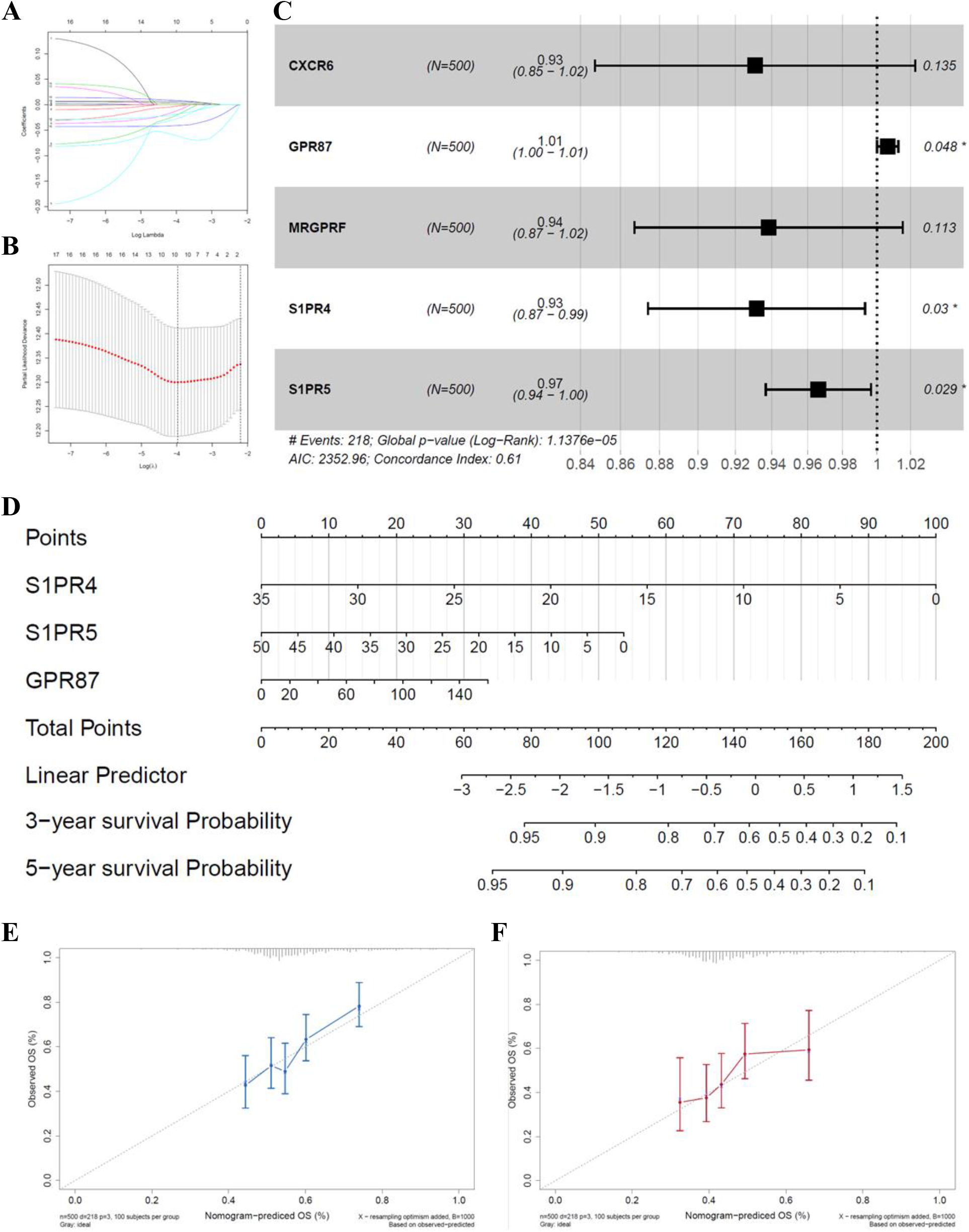
GPR87, S1PR4, and S1PR5 were the potential key GPCRs in HNSCC. **(A, B)** LASSO regression was applied to prevent overfitting. Trajectories of the independent variables were visualized in **(A)**, showing the coefficient of each independent variable at different lambda. The confidence interval at each lambda was visualized in **(B)**. **(C)**Reduced multi-cox regression model was constructed based on the results of LASSO regression (C-Index = 0.61, p < 0.05). Three variables, including GPR87 (Hazard ratio > 1, p = 0.048), S1PR4 (Hazard ratio < 1, p = 0.030), and S1PR5 (Hazard ratio < 1, p = 0.029), were identified as the potential key GPCRs predicting the prognosis of HNSCC. **(D)**According to the reduced multi-cox model in **(C)**, a nomogram was constructed based on GPR87, S1PR4, and S1PR5. **(E, F)** Calibration curves of 3-year survival **(E)** and 5-year survival **(F)** indicated the acceptable accuracy. *GPCR, G protein-coupled receptor; HNSCC, head and neck squamous cell carcinoma; LASSO, Least absolute shrinkage and selection operator. * p < 0*.*05*.

Considering GPR87, S1PR4, and S1PR5 are potential key GPCRs in HNSCC, we further compared these three genes from several perspectives to identify the most important ones. First, all three GPCRs effectively evaluated the prognosis of OS (p < 0.05, **Figure 5A**) and PFS (p < 0.05, **Figure 5B**) in HNSCC patients. Second, only GPR87 (p < 0.05) and S1PR4 (p < 0.05) were identified as DEGs between the two subtypes (**Figure 5B**). Although S1PR5 was one of the featured GPCRs in NMF clustering, it was not differentially expressed between subtypes (**Figure 5B**). Third, we performed another machine learning approach, the RF method, to compare the importance of these three genes in classifying the two molecular subtypes. The results of the RF method identified S1PR4 as the most important GPCR (**Figure 5D**), suggesting that S1PR4 is a key GPCR that determines the abovementioned subtypes. Most patients in subtype C1 belonged to the S1PR4-Low-Expression groups (**Figure S3**). In particular, the correlation analysis indicated that, among the three GPCRs, only S1PR4 was significantly correlated with the proportion and cytotoxicity markers of CD8^+^ T cells (r > 0.4, p < 0.05, **Figure 5E**). Collectively, we identified S1PR4 as a potential key GPCR in HNSCC and selected S1PR4 for further analysis.

**Figure 5.**
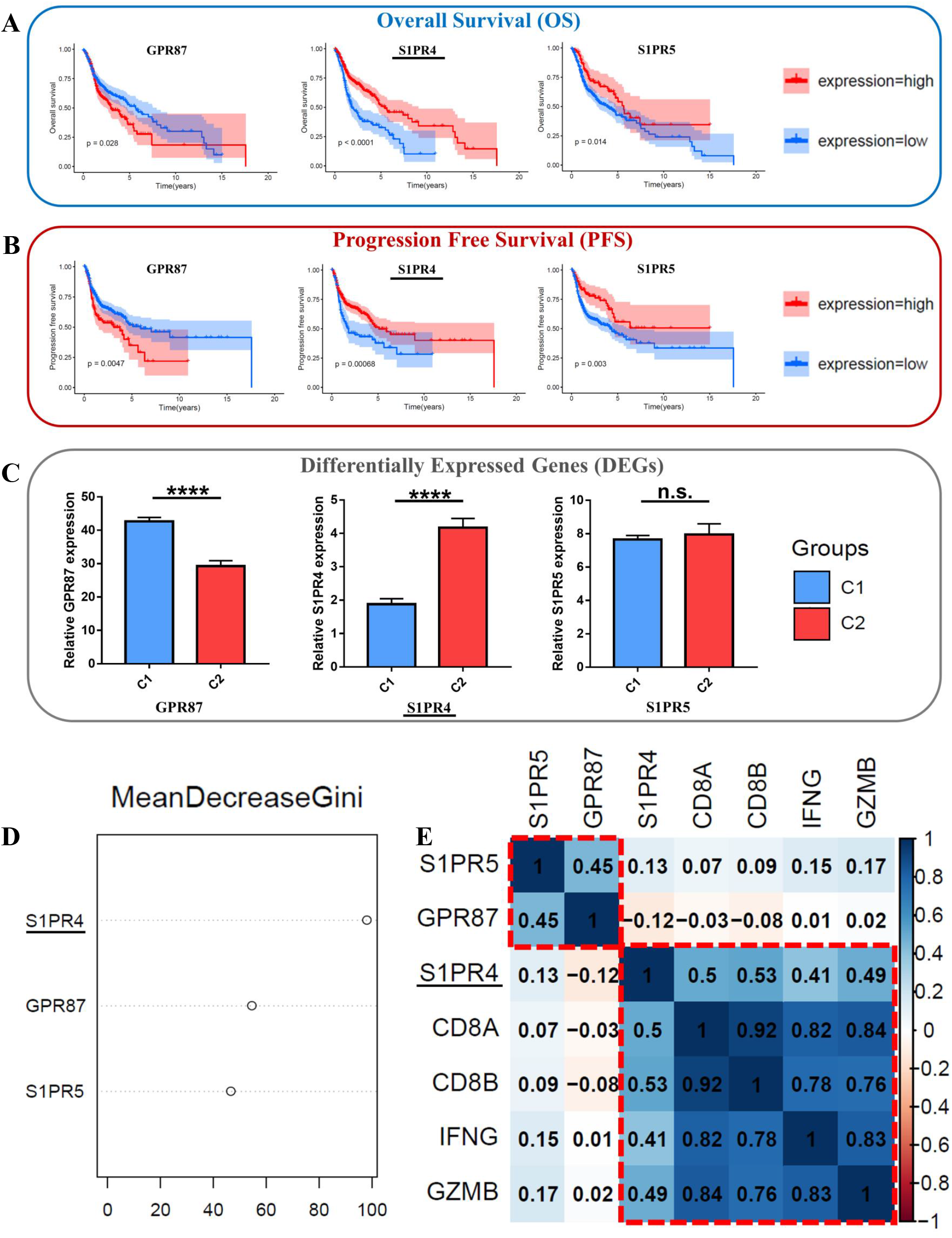
Identification of the Key GPCR S1PR4. **(A)**Kaplan-Meier plots revealed that all of the three GPCRs (GPR87, S1PR4, and S1PR5) could effectively evaluate the prognosis of OS for the HNSCC patients. HNSCC patients with higher S1PR4 (p < 0.0001) or S1PR5 (p < 0.05) expression tended to have better prognosis of OS, while patients with higher GPR87 (p < 0.05) expression tended to have poorer prognosis of OS. **(B)**Kaplan-Meier plots revealed that all of the three GPCRs (GPR87, S1PR4, and S1PR5) could effectively evaluate the prognosis of PFS for the HNSCC patients. HNSCC patients with higher S1PR4 (p < 0.001) or S1PR5 (p < 0.01) expression tended to have better prognosis of PFS, while patients with higher GPR87 (p < 0.01) expression tended to have poorer prognosis of PFS. **(C)**Relative expression of all the three GPCRs (GPR87, S1PR4, and S1PR5) in subtype C1 and C2. GPR87 was significantly downregulated in subtype C2 (p < 0.0001), while the expression of S1PR4 was significantly higher in subtype C2 (p < 0.0001). S1PR5 was not differentially expressed between two subtypes (p > 0.05). **(D)**The RF method was performed to compare the importance of the three GPCRs (GPR87, S1PR4, and S1PR5), revealing that S1PR4 was the most important GPCR in determining the two subtypes. **(E)**Correlation heatmap of the three GPCRs (GPR87, S1PR4, and S1PR5) and the CD8^+^ T cell-related markers. S1PR4 was strongly correlated with the proportion (CD8A and CD8B) and cytotoxicity markers (IFNG and GZMB) of CD8^+^ T cells (r > 0.4). However, both of GPR87 and S1PR4 were weakly correlated with the above CD8^+^ T cell-related markers (r < 0.2). *GPCR, G protein-coupled receptor; OS, overall survival; HNSCC, head and neck squamous cell carcinoma; PFS, progression free survival; RF, Random Forest. **** p < 0*.*0001; ns, not significant*.

### S1PR4 as a Key Indicator for Cytotoxicity and Proportion of CD8^+^ T cells

In aforementioned results, we had identified the correlations between S1PR4 and CD8^+^ T cell-related markers (CD8A, CD8B, IFNG, and GZMB). To confirm these correlations, we used several algorithms to estimate the scores of the cytotoxicity and proportion of CD8^+^ T cells. Consistent with our previous results, S1PR4 was found to be significantly correlated with these scores (p < 0.05, **Figure 6A**), which indicated that S1PR4 might be a key indicator of cytotoxicity and the proportion of CD8^+^ T cells. Therefore, we decided to explore the relationship between S1PR4 and CD8^+^ T cells more comprehensively.

**Figure 6.**
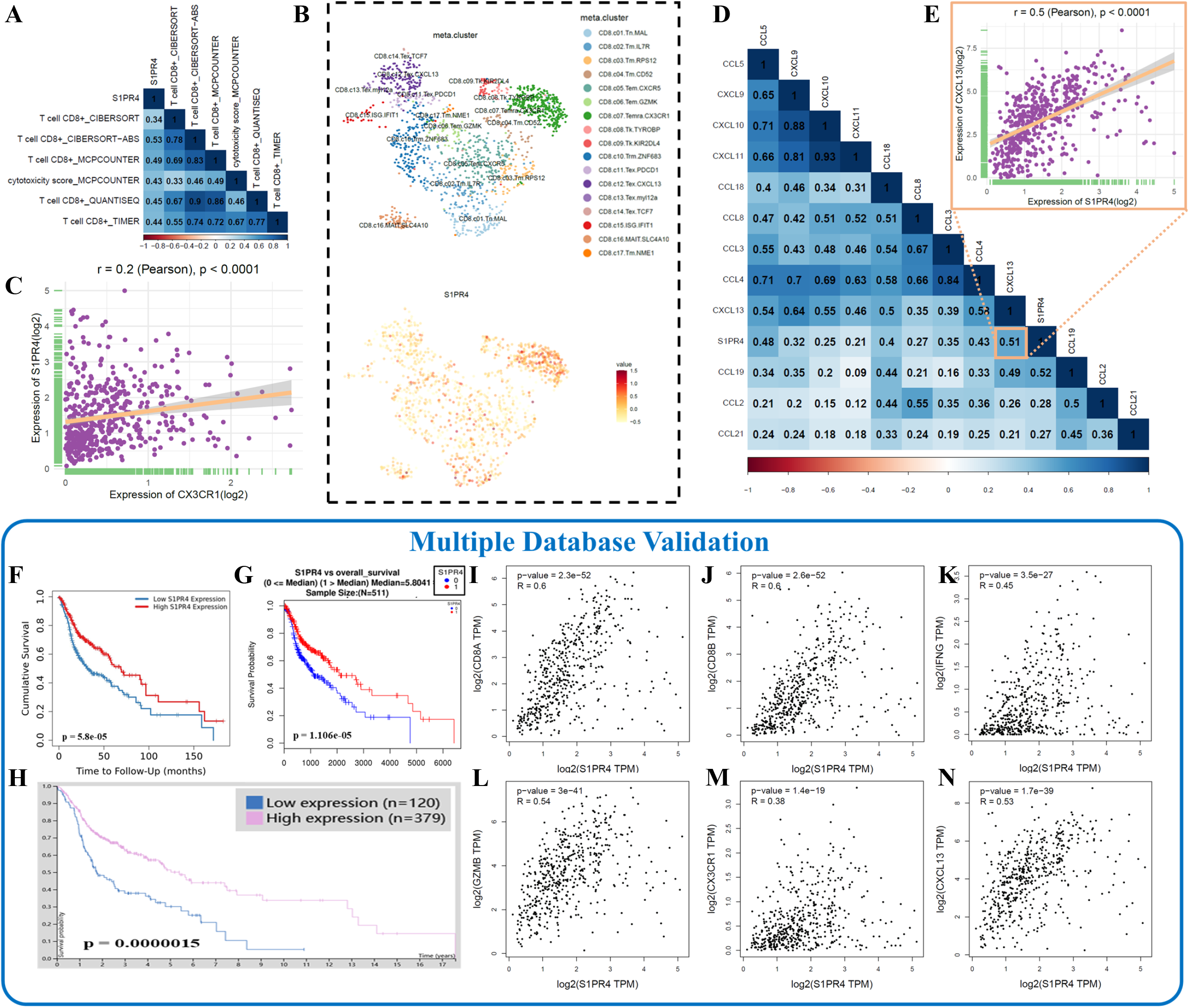
Multiple Database analysis of the relationship between S1PR4 and CD8^+^ T cells. **(A)**Correlation heatmap indicated that S1PR4 was significantly correlated with the algorithm-estimated scores of CD8^+^ T cell cytotoxicity and proportion (p < 0.05). The score of CD8^+^ T cell cytotoxicity was estimated using the MCPcounter algorithm. The scores of CD8^+^ T cell proportion were estimated using five algorithms, including MCPcounter, CIBERSORT, CIBERSORT-ABS, QUANTISEQ, and TIMER. **(B)**S1PR4 was mainly expressed in a subset of CX3CR1^+^CD8^+^ T cells (marked in dark green). **(C)**The dot plot indicated that S1PR4 was significantly correlated with CX3CR1 (r = 0.2, p < 0.0001). **(D)**Correlation heatmap of S1PR4 and the gene signatures of TLS (CCL2, CCL3, CCL4, CCL5, CCL8, CCL18, CCL19, CCL21, CXCL9, CXCL10, CXCL11, and CXCL13). S1PR4 was significantly correlated with all the TLS signatures (p < 0.05). **(E)**The dot plot indicated that S1PR4 was significantly correlated with CXCL13 (r = 0.5, p < 0.0001). **(F-H)** The TIMER database **(F)** indicated that S1PR4 could effectively evaluate the prognosis of HNSCC patients (p < 0.05). Similarly, the prognostic value of S1PR4 was validated using the LinkedOmics database **(G)** and the HumanProteinAtlas database **(H)**. All three databases indicated that patients with higher S1PR4 expression tended to have a better prognosis (p < 0.05). **(I-N)** The Gepia database was used to validate the correlations between S1PR4 and CD8^+^ T cell-related markers, including CD8A **(I)**, CD8B **(J)**, IFNG **(K)**, GZMB **(L)**, and CX3CR1 **(M)**. Similarly, the Gepia database was also used to validate the correlation between S1PR4 and the TLS signature CXCL13 **(N)**. *TLS, tertiary lymphoid structure; head and neck squamous cell carcinoma*.

On the one hand, we used the public single-cell RNA-seq profiles to explore the S1PR4 expression in CD8^+^ T cells. In previous studies, the CX3CR1^+^CD8^+^ T cells were proven to have the highest cytotoxicity score among CD8^+^ T cells[13, 14]. Notably, in our study, S1PR4 was found to be mainly expressed in a subset of CX3CR1^+^CD8^+^ T cells among CD8^+^ T cells (**Figure 6B**). The correlation between S1PR4 and CX3CR1 was also validated using the bulk RNA-seq profiles of TCGA (p < 0.05, **Figure 6C**). On the other hand, recent studies have shown that TLS structure is an important indicator for the cytotoxicity and proportion of CD8^+^ T cells in TIME. And the gene signatures of TLS had been already defined by either a single gene (CXCL13) or multiple genes (CCL2, CCL3, CCL4, CCL5, CCL8, CCL18, CCL19, CCL21, CXCL9, CXCL10, CXCL11, and CXCL13)[36, 37]. Therefore, we also investigated the correlation between TLS and S1PR4 expression. We found that the expression of S1PR4 was significantly correlated with all TLS signatures (p < 0.05, **Figure 6D**), including CXCL13 (r = 0.5, p < 0.05, **Figure 6E**).

### Multiple Database Validation

To prevent the bias from only using a single database, multiple databases were used for validation. The prognostic value of S1PR4 was validated by using the TIMER database (p < 0.05, **Figure 6F**), the LinkedOmics database (p < 0.05, **Figure 6G**), and the HumanProteinAtlas database (p < 0.05, **Figure 6H**), respectively. Moreover, we validated the correlations between S1PR4 and CD8^+^T cell-related markers (CD8A, CD8B, IFNG, GZMB, and CX3CR1) using the Gepia database (**Figures 6I-M**). Similarly, the correlations between S1PR4 and TLS signatures were also validated (**Figure 6N** and **Figure S4**).

### Independent HNSCC Cohort Validation

To validate our bioinformatic findings, an independent cohort from our institution was used. A total of 59 HNSCC patients, with a median follow-up time of 4.17 years, were included in this independent cohort.

First, we explored the correlations among S1PR4, CD8^+^ T cells, and several cytotoxic markers (IFNG and GZMB). Based on IHC staining, the IFNG and GZMB expression was quantified at the protein level (**Figure 7A**). Similarly, the protein expression of CD8 and S1PR4 was quantified in 59 HNSCC patients by IF staining (**Figure 7B**). The heatmap showed the relative expression of these markers (**Figure 7C**). The correlation heatmap revealed that S1PR4 was significantly correlated with CD8 (r= 0.82, p < 0.05), IFNG (r = 0.25, p < 0.05), and GZMB (r = 0.38, p < 0.05), indicating that S1PR4 is essential for both the proportion and cytotoxicity of CD8^+^ T cells (**Figure 7D**).

**Figure 7.**
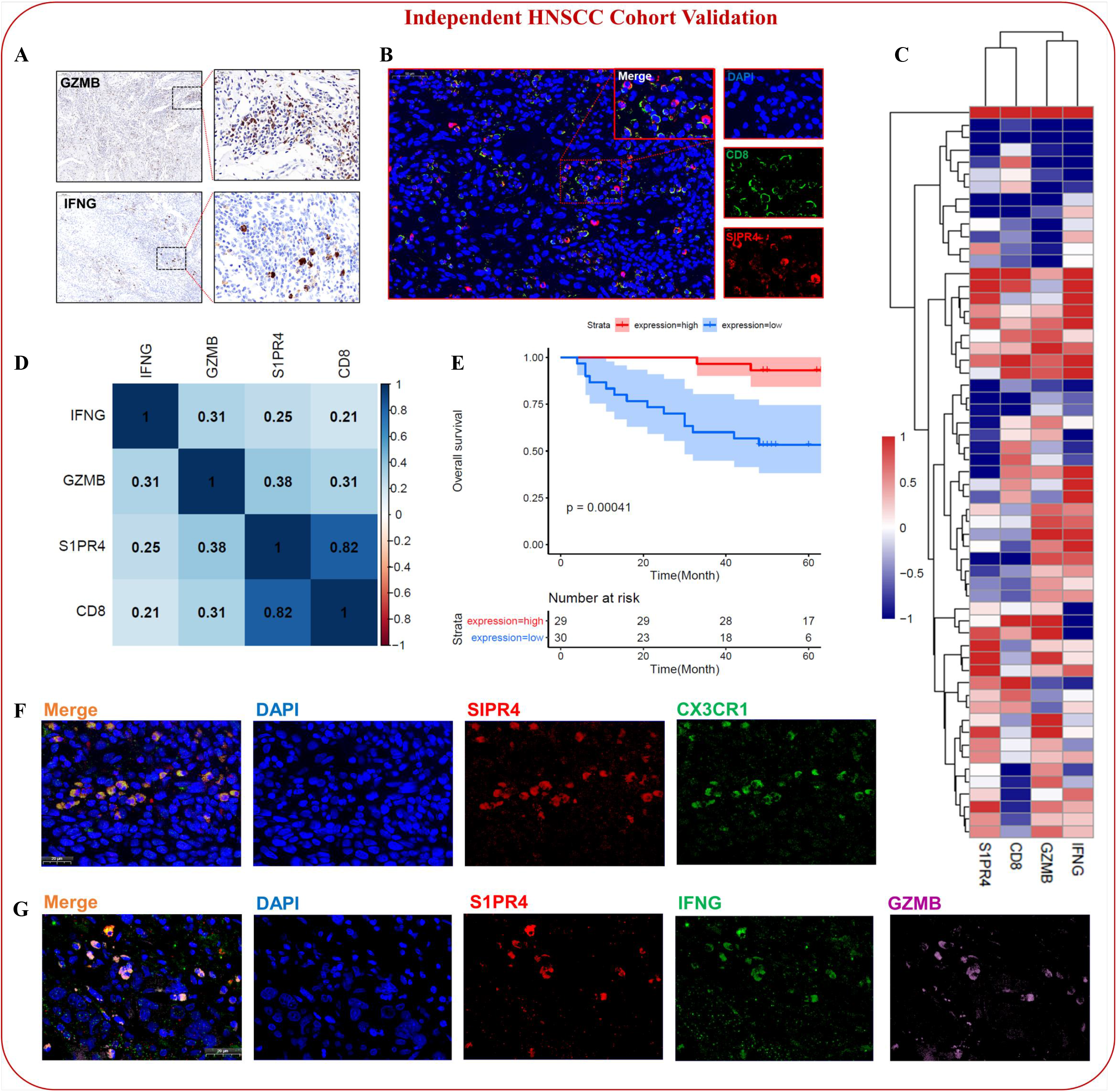
Independent HNSCC Cohort Validation. **(A)**Representative images of GZMB and IFNG IHC staining in the independent HNSCC cohort. **(B**)Representative images of S1PR4 and CD8 IF staining in the independent HNSCC cohort, showing the co-localization between S1PR4 and CD8. **(C)**Heatmap showed the relative protein expression of S1PR4, CD8, GZMB, and IFNG in the independent HNSCC cohort. **(D)**Correlation heatmap of S1PR4, CD8, GZMB, and IFNG. At the protein level, S1PR4 was found to have the strongest correlation with CD8 (r = 0.82, p < 0.05). The S1PR4 was also found to be significantly correlated with both IFNG (r = 0.25, p < 0.05) and GZMB (r = 0.38, p < 0.05). **(E)**The Kaplan-Meier plot revealed that, at the protein level, the HNSCC patients with higher S1PR4 expression tended to have a better prognosis (p = 0.00041). **(F)**Representative images of the co-localization between S1PR4 and the cytotoxic markers of CD8^+^ T cells (IFNG and GZMB). **(G)**Representative images of the co-localization between S1PR4 and CX3CR1. *HNSC, Head and neck squamous cell carcinoma*.

Then, we performed K-M analysis, confirming that HNSCC patients with higher S1PR4 protein expression tended to have a better OS (p < 0.05, **Figure 7E**), which was consistent with our bioinformatics findings.

After statistically confirming the correlations among S1PR4, CD8^+^ T cells, and the cytotoxic markers, we further explored the spatial distributions among S1PR4 protein, CD8^+^ T cells, and the cytotoxic markers. Based on IF staining experiments, the colocalisation of S1PR4 with CD8^+^ T cells was confirmed (**Figure 7B**), suggesting that S1PR4 might be essential for CD8^+^ T cells. In addition, considering that we had found that S1PR4 was mainly expressed in a subset of CX3CR1^+^CD8^+^ T cells, we further validated the co-localisation between S1PR4 and CX3CR1 (**Figure 7F**). According to the aforementioned literature research results, the CX3CR1^+^CD8^+^ T cells had already been identified as the most cytotoxic subset of CD8^+^ T cells. Therefore, we also confirmed the co-localisation of S1PR4 and the important cytotoxic markers of CD8^+^ T cells (IFNG and GZMB, **Figure 7G**). Collectively, S1PR4 was found to be expressed in the most cytotoxic subset of CD8^+^ T cells (CX3CR1^+^CD8^+^ T cells), and was also found to have co-localisations with important cytotoxic markers (IFNG and GZMB) of CD8^+^ T cells. In short, S1PR4 might play an important role in the CX3CR1^+^CD8^+^ T cell-mediated tumour killing, which is essential for anti-tumour immune regulation in TME.

### Evaluation of the Function of S1PR4 in Cancer Immunotherapy

To further evaluate S1PR4’s function in cancer immunotherapy, we generated S1PR4-downregulated and S1PR4-upregulated PSMA-CAR-T cells using an S1PR4 inhibitor (CYM50358) and a S1PR4 agonist (CYM50308), respectively. **Figure 8A** shows the chemical structures of CYM50358 and CYM50308. The cytotoxicity of CAR-T cells was significantly repressed in the S1PR4-downregulated group, and enhanced in the S1PR4-upregulated group (**Figure 8B**). Moreover, we also measured the expression of GZMB in PSMA-CAR-T cells using flow cytometry after co-incubation with PC3 cells (**Figure 8C**). The S1PR4 inhibitor (CYM50358) significantly decreased GZMB release, whereas the S1PR4 agonist (CYM50308) increased GZMB release (**Figure 8D**). Similarly, the S1PR4 agonist (CYM50308) could considerably increase the release of IFNG (**Figure 8E**). Collectively, the upregulation of S1PR4 enhanced the cytotoxicity of CAR-T cells and promoted the release of important cytotoxic markers (GZMB and IFNG).

**Figure 8.**
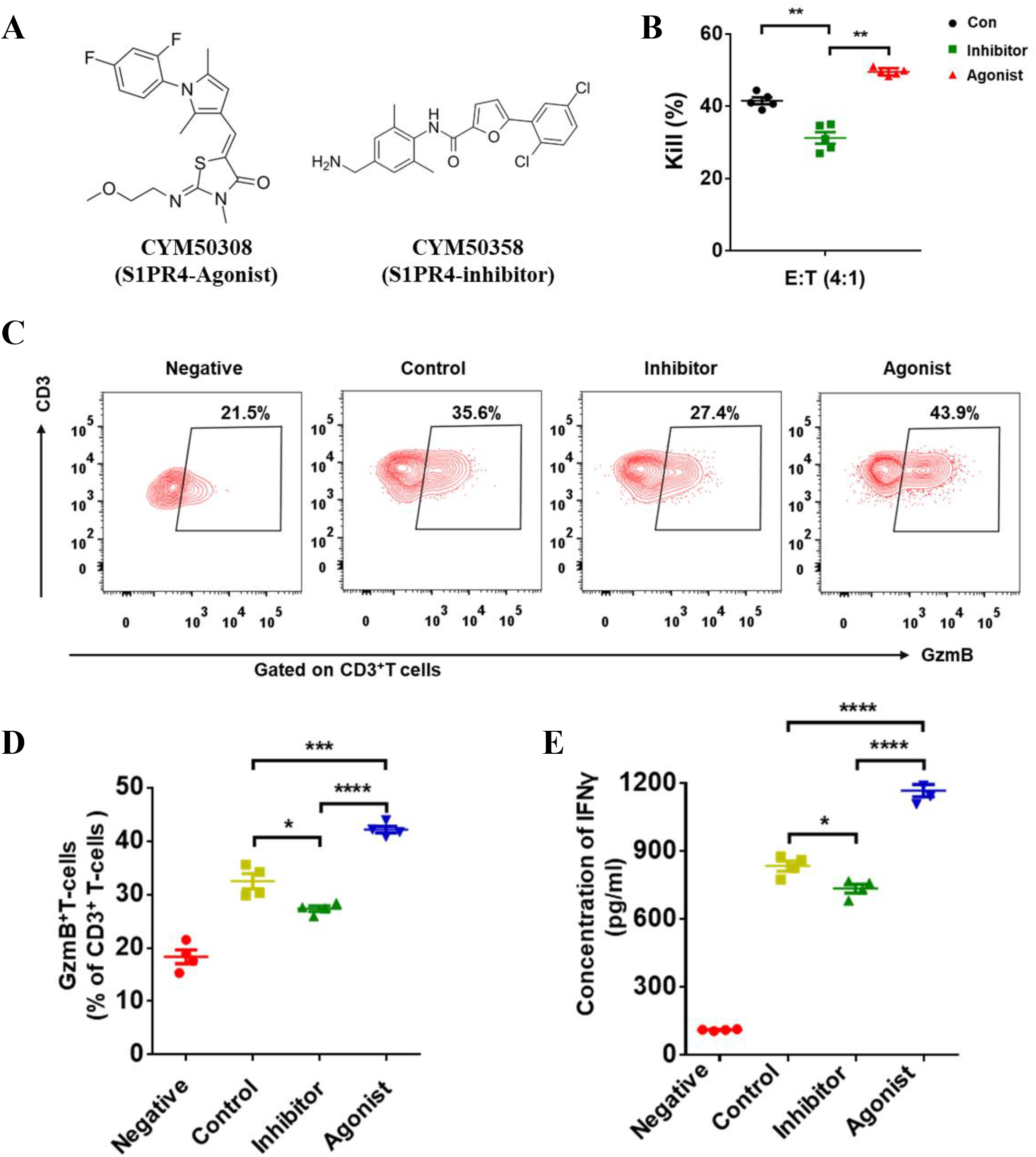
Evaluation of the Function of S1PR4 in Cancer Immunotherapy. **(A)**Chemical structures of S1PR4 agonist (CYM50308, left) and S1PR4 inhibitor (CYM50358, right). **(B)**PSMA-CAR-T cells were pretreated with DMSO, S1PR4-inhibitor (CYM50358; 100 nM), and S1PR4-agonist (CYM50308; 100 nM) for 2 h. PSMA-CAR-T against the PC3 tumour cells at an effector-to-target ratio of 4:1 for 12 h. **(C)**PSMA-CAR-T cells were pretreated with DMSO, S1PR4-inhibitor (CYM50358; 100 nM) and S1PR4-agonist (CYM50308; 100 nM) for 2 h. PSMA-CAR-T cells were co-cultured without PC3-PSMA^+^ target tumour cells (negative control) or with 1 × 10^6^ PC3-PSMA^+^ target tumour cells at an effector-to-target ratio of 2:1 in triplicates in 6-well plates (Corning) in a total volume of 2ml of cell media. **(D)**The GZMB expression of PSMA-CAR-T was measured using flow cytometry after co-incubating with PC3 cells at a 2:1 ratio for 16 h. **(E)**Concentration of IFNG in the cultured medium was measured using ELISA. *CAR-T, Chimeric antigen receptor T cell. * p < 0*.*05; ** p < 0*.*01; *** p < 0*.*001; **** p < 0*.*0001*.

## Discussion

The GPCR superfamily is one of the most important families of human proteins. Regulating a wide range of physiological processes, it is known to be essential in the progression of multiple tumour types, including HNSCC. Additionally, as cell surface proteins, GPCRs have broad prospects in the field of drug development. However, currently, the biological functions of many GPCRs remain unclear, and further study of their involvement in tumour progression is still warranted. Therefore, this study aimed to explore the key GPCRs involved in HNSCC progression and provide a potential GPCR-targeted therapeutic option for tumour treatment.

In this study, we adopted the strategy of “classification before analysis”. First, from a statistical perspective, we classified the HNSCC patients into two molecular subtypes only based on their gene expression patterns. Then, from clinical and biological perspectives, we comprehensively compared the two subtypes in terms of prognosis, gene expression, pathway expression, TIME, and immune cell infiltration. By exploring the featured genes between the two subtypes, we identified S1PR4 as a potential key GPCR in HNSCC.

The GPCR superfamily is defined based on the characteristic of the seven-transmembrane structure, and the biological function of each GPCR varies. When using the strategy of “classification before analysis”, we did not limit the biological differences in the initial classification of molecular subtypes. Therefore, in the process of classification, we could only determine that there were statistical differences in HNSCC patients’ gene expression patterns, but could not determine the differences in specific biological functions between the two molecular subtypes. For comparison, previous similar studies often obtained the gene list of specific biological functions or pathways in advance, and then classified the molecular subtypes based on these selected genes. Compared with this strategy of “determining biological functions before classification”, our strategy did not anticipate biological differences between subtypes, which made our strategy relatively more objective.

Using this strategy, our study identified S1PR4 as a key GPCR in HNSCC. The GPCR S1PR4 belongs to the endothelial differentiation G protein-coupled (EDG) receptors, which ligands includes lysophosphingolipids and sphingolipids[38]. In the past decades, EDG receptors have been shown to be participate in signal transduction in multiple cell types, indicating their importance in physiological and pathological regulation. As a member of EDG receptors, S1PR4 binds sphingosine-1-phosphate (S1P) as the ligand, and has been found to play a role in immune regulation.

In previous studies, S1PR4 has been reported to regulate the migration and proliferation of immune cells, especially T cells. For instance, Graeler et al. and Xiong et al. showed that S1PR4 plays an important role in promoting T cell migration[39, 40]. And Schulze et al. demonstrated that deletion of S1PR4 would affect S1P-induced CD8^+^ T cell migration[41]. Meanwhile, Olesch et al. showed that the ablation of S1PR4 promotes the expansion of CD8^+^ T cells[42]. In short, S1PR4 could not only promote CD8^+^ T cell migration, but also inhibit CD8^+^ T cell expansion. Therefore, the relationship between S1PR4 and the proportion of tumour-infiltrating CD8^+^ T cells depends on the joint action of the above two regulatory mechanisms, which have opposite effects on the regulation of CD8^+^ T cell proportion. Notably, a previous study presumed that S1PR4 might play a minor role in lymphocyte migration rather than expansion[43]. And a recent study further endorsed the possibility that S1PR4 could drive CXCR4, affecting lymphocyte differentiation (for example, expansion) rather than lymphocyte trafficking[44]. However, the relationship between S1PR4 and the proportion of CD8^+^ T cells in HNSCC remains largely unknown. Our study identified that, at the tissue level in HNSCC samples, S1PR4 is positively correlated with the proportion of tumour-infiltrating CD8^+^ T cells, which contributes to anti-tumour immunity. Considering that HNSCC is a special tumour type with relatively high lymphocyte infiltration, we hypothesised that S1PR4 might play a major role in CD8^+^ T cell migration rather than expansion in HNSCC patients.

Moreover, we also identified that S1PR4 was positively correlated with the cytotoxicity of CD8^+^ T cells in HNSCC, and further found that S1PR4 was mainly expressed in a subset of CX3CR1^+^CD8^+^ T cells. Interestingly, in the literature, the CX3CR^+^CD8^+^ T cells were identified as a high migration subset, and were further reported to have the highest cytotoxicity score among CD8^+^ T cells[13, 14]. On the other hand, in recent years, many studies have confirmed that the TLS is an important effector of anti-tumour immunity, and could promote the cytotoxicity of CD8^+^ T cells[36, 45, 46]. In our study, S1PR4 was also found to be positively correlated with gene signatures of TLSs in HNSCC. Collectively, our findings are in accordance with the above literatures, suggesting that S1PR4, which is mainly expressed in CX3CR1^+^CD8^+^ T cells, might be essential for the cytotoxic functions of tumour-infiltrating CD8^+^ T cells.

Several limitations in this study need to be addressed. First, this study only focused on the relationship between S1PR4 and CD8^+^ T cells, and did not further analyse other types of immune cells. Second, most of the findings in this study were based on bioinformatics, and did not further investigate the underlying mechanisms. In future studies, the biological mechanisms of S1PR4 regulating CD8^+^ T cells’ cytotoxicity should be further investigated experimentally. Despite the abovementioned limitations, this study is the first to infer that S1PR4 might be a key GPCR in HNSCC. We found that S1PR4 could effectively evaluate the prognosis of both OS and PFS in patients with HNSCC. And we also found that S1PR4 was mainly expressed in CX3CR1^+^CD8^+^ T cells, and was positively correlated with both the proportion and cytotoxicity of CD8^+^ T cells in HNSCC. The above findings were validated using an independent cohort from our institution, suggesting that S1PR4 might be a potential therapeutic target for HNSCC treatment.

In summary, this study proposed a strategy of “classification before analysis”, and identified S1PR4 as a key GPCR for HNSCC patients. We found that S1PR4 was a key GPCR affecting HNSCC patient prognosis, and further constructed a prognosis-predicting nomogram based on three variables, including S1PR4. We also found that S1PR4 was mainly expressed in CX3CR1^+^CD8^+^ T cells, and was positively correlated with the proportion and cytotoxicity of CD8^+^ T cells. Moreover, S1PR4 enhanced CAR-T cell cytotoxicity and promoted the release of important cytotoxic markers (GZMB and IFNG). Our findings contribute to the knowledge of S1PR4 in anti-tumour immunity, and might provide a potential GPCR-targeted therapeutic option for future HNSCC treatment.

## Competing interests

The authors declare no competing interests.

## Author Contributions

Conception/design: CH.

Literature research: CH, FZ, NW, QH

Data acquisition: CH, FZ, NW

Data analysis/interpretation: CH, FZ, NW

Experimental studies: CH, FZ, NW

Writing, original draft: CH, QH.

Writing, review and editing: CH, QH.

All authors read and approved the submitted version of manuscript.

### Acknowledgements

The authors thank Yue Tian (East China Normal University, Shanghai, China) and Runzhi Huang (Tongji University, Shanghai, China) for generously sharing their experience.

## Data Accessibility

Publicly available datasets were analyzed in this study. Gene expression profiles of HNSCC patients could be acquired from UCSC Xena (https://xenabrowser.net/). Further inquiries could be directed to the corresponding authors.

## Ethics Statement

This study was reviewed and approved by Shanghai Ninth People’s Hospital, Shanghai, China. (SH9H-2022-T189-1).

## Supplementary Material

**Figure S1** | **Consensus map of NMF clustering (rank k was set from 2 to 10)**.

**Figure S2** | **Kaplan-Meier plots indicated that all 17 GPCRs showed prognostic values of HNSCC in the OS of patients**.

**Figure S3** | **Most patients of subtype C1 belonged to the S1PR4-Low-Expression groups**.

**Figure S4** | **The Gepia database was used to validate the correlations between S1PR4 and the gene signatures of TLS (CCL2, CCL3, CCL4, CCL5, CCL8, CCL18, CCL19, CCL21, CXCL9, CXCL10, CXCL11, and CXCL13)**.

## Notes

### Competing Interest Statement

The authors have declared no competing interest.

### Summary of Updates

We have revised our analytical approach and re conducted data analysis, adding experimental validation and clinical sample validation.

## References

1. Lappano, R. and M. Maggiolini, G protein-coupled receptors: novel targets for drug discovery in cancer. Nat Rev Drug Discov, 2011. 10(1): p. 47–60.

2. Lappano, R. and M. Maggiolini, GPCRs and cancer. Acta Pharmacol Sin, 2012. 33(3): p. 351–62.

3. Voisin, T., et al., Aberrant expression of OX1 receptors for orexins in colon cancers and liver metastases: an openable gate to apoptosis. Cancer Res, 2011. 71(9): p. 3341–51.

4. Jassal, B., et al., The systematic annotation of the three main GPCR families in Reactome. Database (Oxford), 2010. 2010: p. baq018.

5. Miyauchi, S., et al., Immune Modulation of Head and Neck Squamous Cell Carcinoma and the Tumor Microenvironment by Conventional Therapeutics. Clin Cancer Res, 2019. 25(14): p. 4211–4223.

6. Solomon, B., R.J. Young, and D. Rischin, Head and neck squamous cell carcinoma: Genomics and emerging biomarkers for immunomodulatory cancer treatments. Semin Cancer Biol, 2018. 52(Pt 2): p. 228–240.

7. Bhat, A.A., et al., Tumor microenvironment: an evil nexus promoting aggressive head and neck squamous cell carcinoma and avenue for targeted therapy. Signal Transduct Target Ther, 2021. 6(1): p. 12.

8. Elmusrati, A., J. Wang, and C.Y. Wang, Tumor microenvironment and immune evasion in head and neck squamous cell carcinoma. Int J Oral Sci, 2021. 13(1): p. 24.

9. Johnson, D.E., et al., Head and neck squamous cell carcinoma. Nat Rev Dis Primers, 2020. 6(1): p. 92.

10. Hanakawa, H., et al., Regulatory T-cell infiltration in tongue squamous cell carcinoma. Acta Otolaryngol, 2014. 134(8): p. 859–64.

11. Seiwert, T.Y. and E.E. Cohen, Targeting angiogenesis in head and neck cancer. Semin Oncol, 2008. 35(3): p. 274–85.

12. Sasidharan Nair, V. and E. Elkord, Immune checkpoint inhibitors in cancer therapy: a focus on T-regulatory cells. Immunol Cell Biol, 2018. 96(1): p. 21–33.

13. Zheng, L., et al., Pan-cancer single-cell landscape of tumor-infiltrating T cells. Science, 2021. 374(6574): p. abe6474.

14. Liu, Y., et al., Tumour heterogeneity and intercellular networks of nasopharyngeal carcinoma at single cell resolution. Nat Commun, 2021. 12(1): p. 741.

15. Pucher, B.M., O.A. Zeleznik, and G.G. Thallinger, Comparison and evaluation of integrative methods for the analysis of multilevel omics data: a study based on simulated and experimental cancer data. Brief Bioinform, 2019. 20(2): p. 671–681.

16. Brunet, J.P., et al., Metagenes and molecular pattern discovery using matrix factorization. Proc Natl Acad Sci U S A, 2004. 101(12): p. 4164–9.

17. Gaujoux, R. and C. Seoighe, A flexible R package for nonnegative matrix factorization. BMC Bioinformatics, 2010. 11: p. 367.

18. Robinson, M.D., D.J. McCarthy, and G.K. Smyth, edgeR: a Bioconductor package for differential expression analysis of digital gene expression data. Bioinformatics, 2010. 26(1): p. 139–40.

19. Subramanian, A., et al., Gene set enrichment analysis: a knowledge-based approach for interpreting genome-wide expression profiles. Proc Natl Acad Sci U S A, 2005. 102(43): p. 15545–50.

20. Yu, G., et al., clusterProfiler: an R package for comparing biological themes among gene clusters. OMICS, 2012. 16(5): p. 284–7.

21. Wu, T., et al., clusterProfiler 4.0: A universal enrichment tool for interpreting omics data. Innovation (Camb), 2021. 2(3): p. 100141.

22. Becht, E., et al., Estimating the population abundance of tissue-infiltrating immune and stromal cell populations using gene expression. Genome Biol, 2016. 17(1): p. 218.

23. Newman, A.M., et al., Robust enumeration of cell subsets from tissue expression profiles. Nat Methods, 2015. 12(5): p. 453–7.

24. Newman, A.M., et al., Determining cell type abundance and expression from bulk tissues with digital cytometry. Nat Biotechnol, 2019. 37(7): p. 773–782.

25. Plattner, C., F. Finotello, and D. Rieder, Deconvoluting tumor-infiltrating immune cells from RNA-seq data using quanTIseq. Methods Enzymol, 2020. 636: p. 261–285.

26. Li, B., et al., Comprehensive analyses of tumor immunity: implications for cancer immunotherapy. Genome Biol, 2016. 17(1): p. 174.

27. Ritchie, M.E., et al., limma powers differential expression analyses for RNA-sequencing and microarray studies. Nucleic Acids Res, 2015. 43(7): p. e47.

28. Friedman, J., T. Hastie, and R. Tibshirani, Regularization Paths for Generalized Linear Models via Coordinate Descent. J Stat Softw, 2010. 33(1): p. 1–22.

29. Kazius, J., et al., GPCR NaVa database: natural variants in human G protein-coupled receptors. Hum Mutat, 2008. 29(1): p. 39–44.

30. Li, T., et al., TIMER: A Web Server for Comprehensive Analysis of Tumor-Infiltrating Immune Cells. Cancer Res, 2017. 77(21): p. e108–e110.

31. Li, T., et al., TIMER2.0 for analysis of tumor-infiltrating immune cells. Nucleic Acids Res, 2020. 48(W1): p. W509–W514.

32. Vasaikar, S.V., et al., LinkedOmics: analyzing multi-omics data within and across 32 cancer types. Nucleic Acids Res, 2018. 46(D1): p. D956–D963.

33. Tang, Z., et al., GEPIA: a web server for cancer and normal gene expression profiling and interactive analyses. Nucleic Acids Res, 2017. 45(W1): p. W98–W102.

34. Devarajan, K., Nonnegative matrix factorization: an analytical and interpretive tool in computational biology. PLoS Comput Biol, 2008. 4(7): p. e1000029.

35. Huang, J., et al., Evaluation of gene-drug common module identification methods using pharmacogenomics data. Brief Bioinform, 2021. 22(3).

36. Petitprez, F., et al., B cells are associated with survival and immunotherapy response in sarcoma. Nature, 2020. 577(7791): p. 556–560.

37. Coppola, D., et al., Unique ectopic lymph node-like structures present in human primary colorectal carcinoma are identified by immune gene array profiling. Am J Pathol, 2011. 179(1): p. 37–45.

38. Graler, M.H., et al., The sphingosine 1-phosphate receptor S1P4 regulates cell shape and motility via coupling to Gi and G12/13. J Cell Biochem, 2003. 89(3): p. 507–19.

39. Graeler, M., G. Shankar, and E.J. Goetzl, Cutting edge: suppression of T cell chemotaxis by sphingosine 1-phosphate. J Immunol, 2002. 169(8): p. 4084–7.

40. Xiong, Y., et al., CD4 T cell sphingosine 1-phosphate receptor (S1PR)1 and S1PR4 and endothelial S1PR2 regulate afferent lymphatic migration. Sci Immunol, 2019. 4(33).

41. Schulze, T., et al., Sphingosine-1-phospate receptor 4 (S1P(4)) deficiency profoundly affects dendritic cell function and TH17-cell differentiation in a murine model. FASEB J, 2011. 25(11): p. 4024–36.

42. Olesch, C., et al., S1PR4 ablation reduces tumor growth and improves chemotherapy via CD8+ T cell expansion. J Clin Invest, 2020. 130(10): p. 5461–5476.

43. Wang, W., M.H. Graeler, and E.J. Goetzl, Type 4 sphingosine 1-phosphate G protein-coupled receptor (S1P4) transduces S1P effects on T cell proliferation and cytokine secretion without signaling migration. FASEB J, 2005. 19(12): p. 1731–3.

44. Burkard, T., et al., Enhanced CXCR4 Expression of Human CD8(Low) T Lymphocytes Is Driven by S1P(4). Front Immunol, 2021. 12: p. 668884.

45. Helmink, B.A., et al., B cells and tertiary lymphoid structures promote immunotherapy response. Nature, 2020. 577(7791): p. 549–555.

46. Cabrita, R., et al., Tertiary lymphoid structures improve immunotherapy and survival in melanoma. Nature, 2020. 577(7791): p. 561–565.

