## Supplementary figures and images for "Identification of S1PR4 as an immune modulator for favourable prognosis in HNSCC through unbiased machine learning"

### Supplementary Figures S1-S4

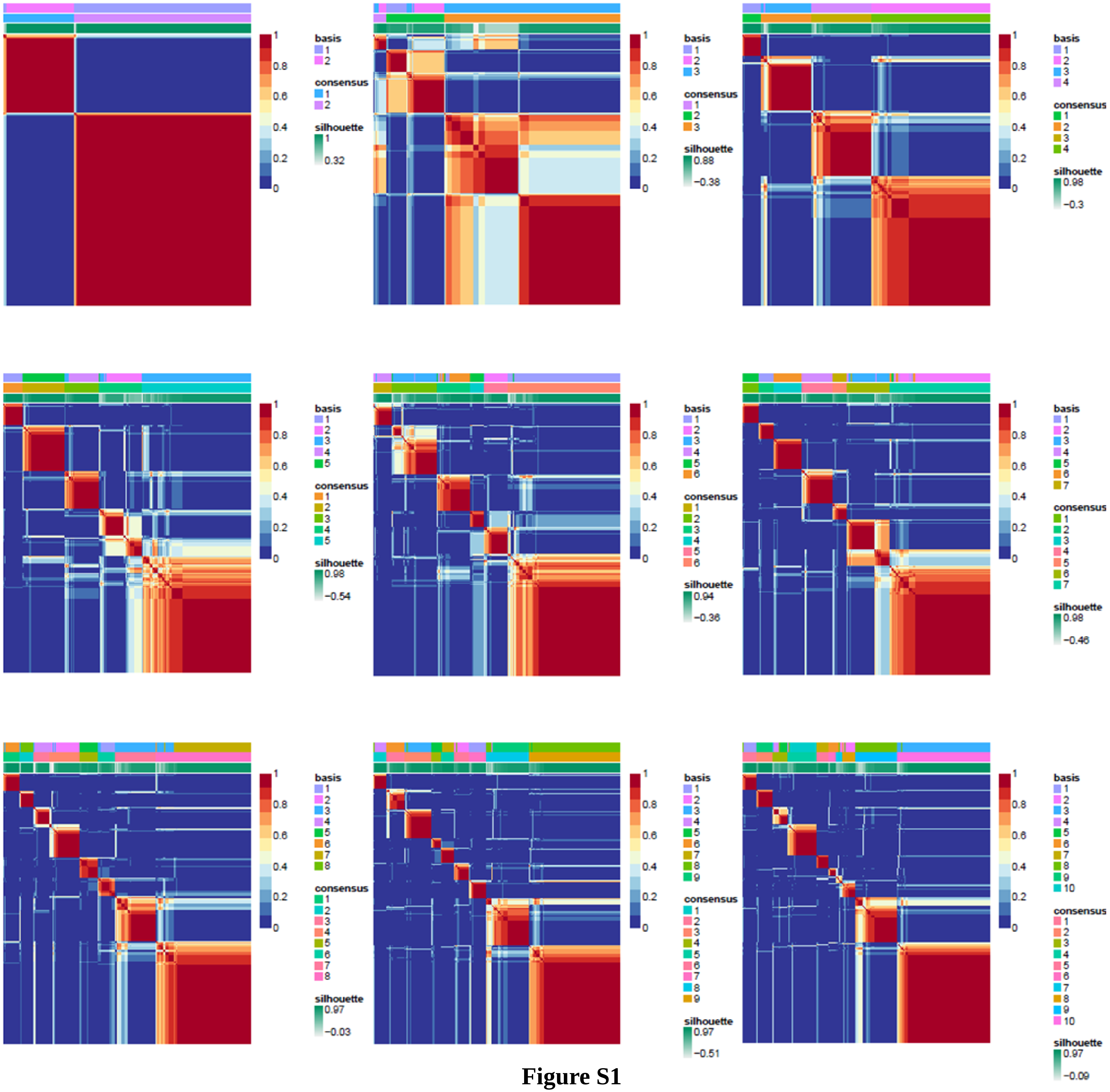

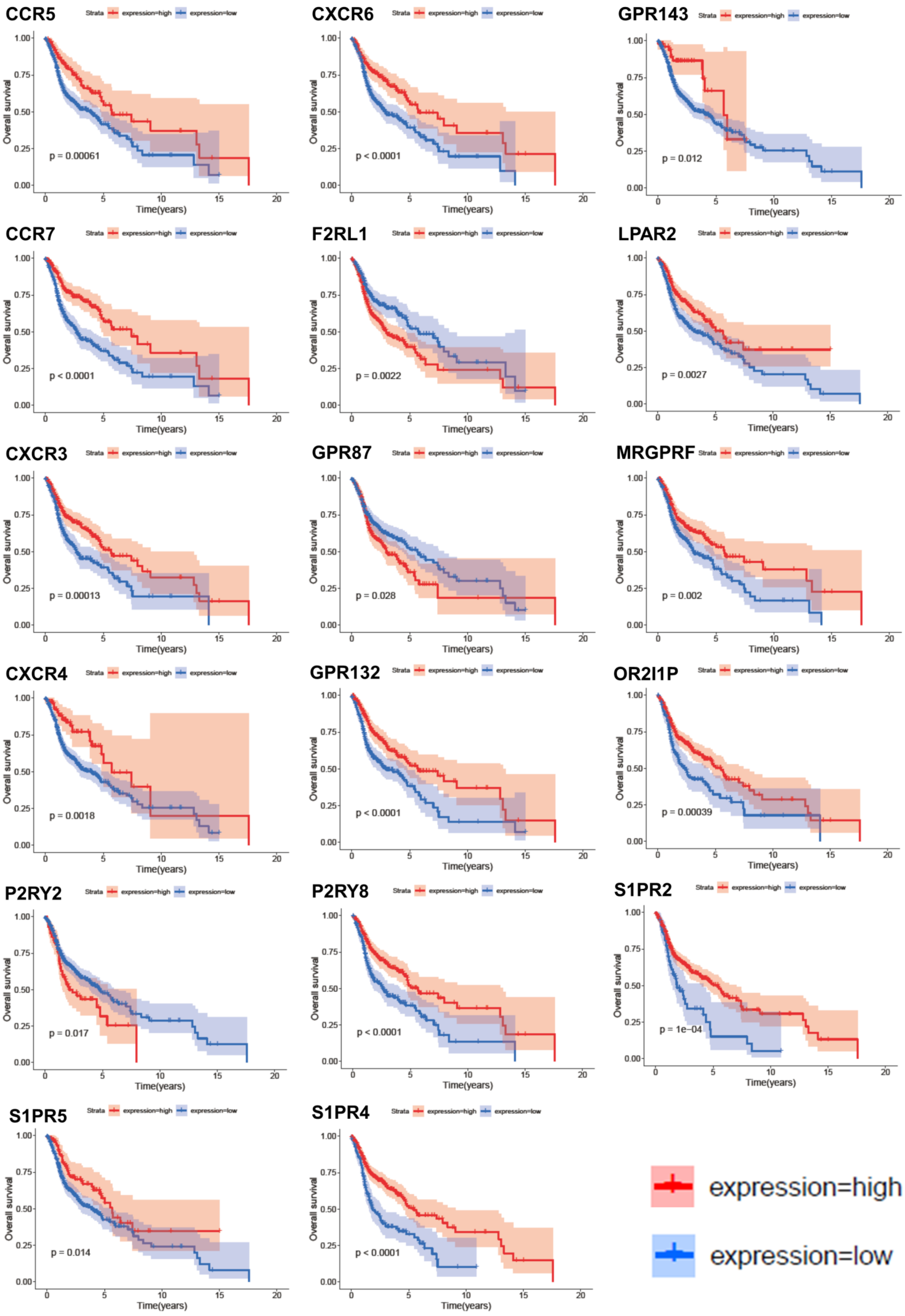

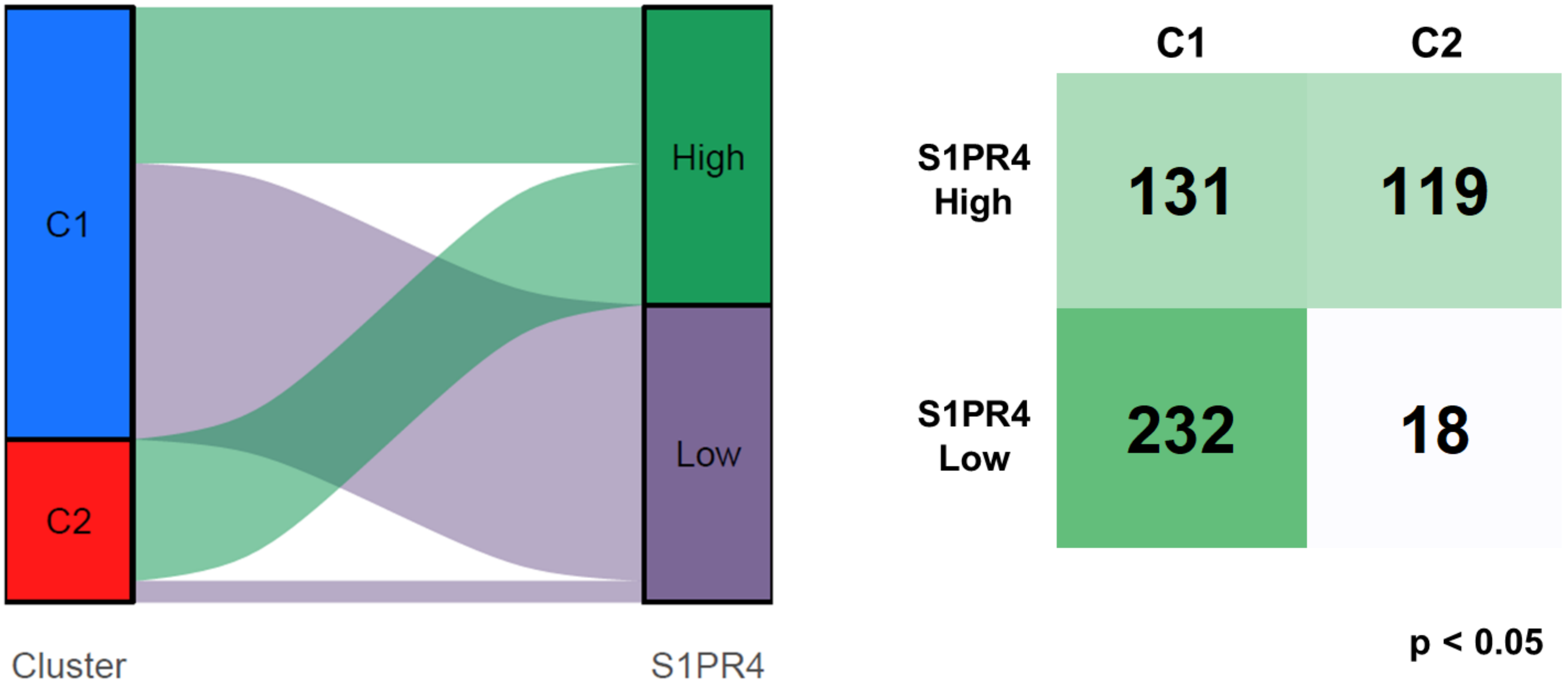

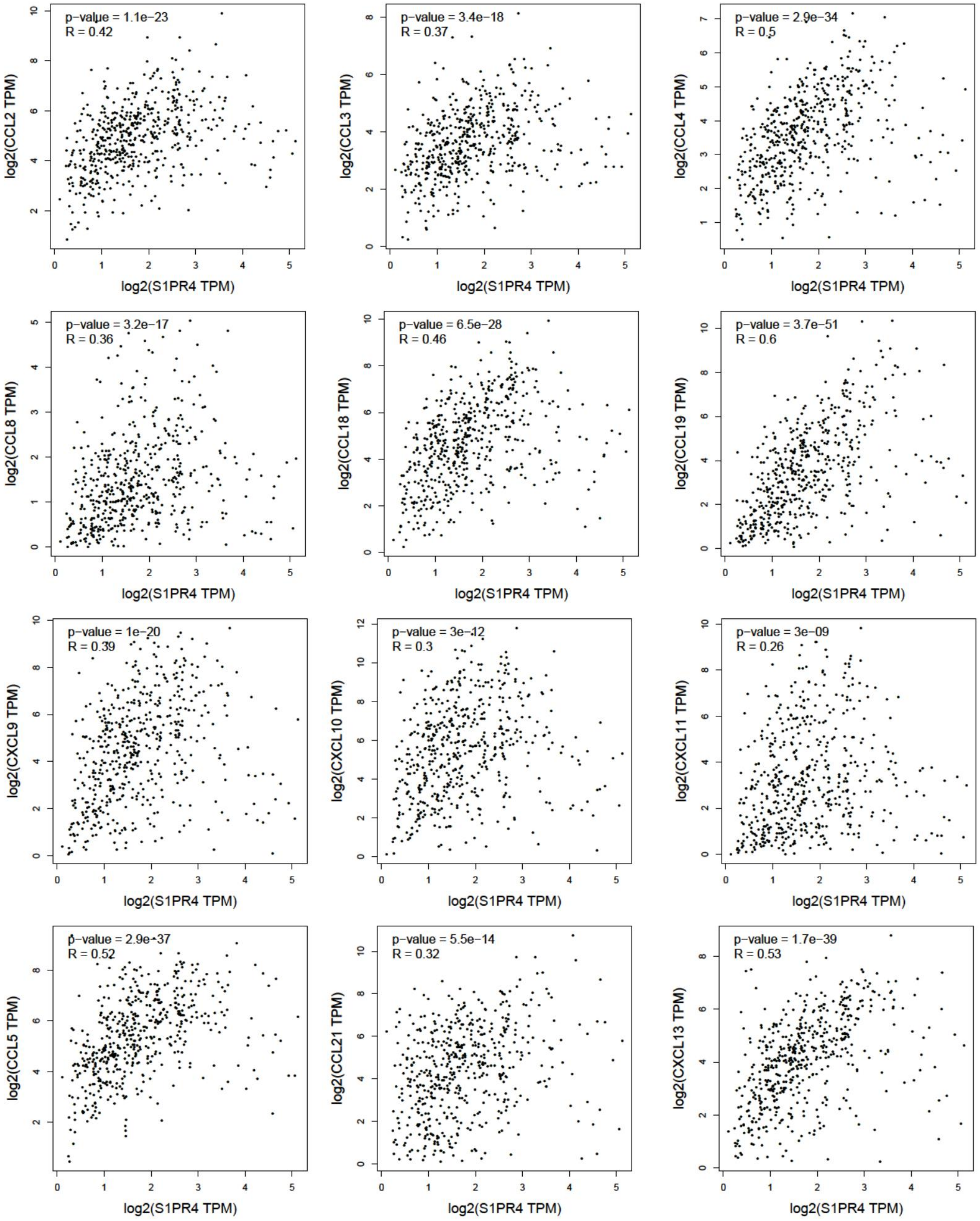
